# Omicron-specific mRNA vaccine induced potent neutralizing antibody against Omicron but not other SARS-CoV-2 variants

**DOI:** 10.1101/2022.01.31.478406

**Authors:** I-Jung Lee, Cheng-Pu Sun, Ping-Yi Wu, Yu-Hua Lan, I-Hsuan Wang, Wen-Chun Liu, Sheng-Che Tseng, Szu-I Tsung, Yu-Chi Chou, Monika Kumari, Yu-Wei Chang, Hui-Feng Chen, Yin-Shiou Lin, Tsung-Yen Chen, Chi-Wen Chiu, Chung-Hsuan Hsieh, Cheng-Ying Chuang, Chih-Chao Lin, Chao-Min Cheng, Hsiu-Ting Lin, Wan-Yu Chen, Po-Cheng Chiang, Chong-Chou Lee, James C. Liao, Han-Chung Wu, Mi-Hua Tao

**Affiliations:** Institute of Biomedical Sciences, Academia Sinica, Taipei, Taiwan; Graduate Institute of Microbiology, College of Medicine, National Taiwan University, Taipei, Taiwan; Biomedical Translation Research Center, Academia Sinica, Taipei, Taiwan; Institute of Cellular and Organismic Biology, Academia Sinica, Taipei, Taiwan; Department of Clinical Laboratory Science and Medical Biotechnology, National Taiwan University, Taipei, Taiwan; Institute of Biological Chemistry, Academia Sinica, Taipei, Taiwan

## Abstract

The emerging SARS-CoV-2 variants of concern (VOC) harbor mutations associated with increasing transmission and immune escape, hence undermine the effectiveness of current COVID-19 vaccines. In late November of 2021, the Omicron (B.1.1.529) variant was identified in South Africa and rapidly spread across the globe. It was shown to exhibit significant resistance to neutralization by serum not only from convalescent patients, but also from individuals receiving currently used COVID-19 vaccines with multiple booster shots. Therefore, there is an urgent need to develop next generation vaccines against VOCs like Omicron. In this study, we develop a panel of mRNA-LNP-based vaccines using the receptor binding domain (RBD) of Omicron and Delta variants, which are dominant in the current wave of COVID-19. In addition to the Omicron- and Delta-specific vaccines, the panel also includes a “Hybrid” vaccine that uses the RBD containing all 16 point-mutations shown in Omicron and Delta RBD, as well as a bivalent vaccine composed of both Omicron and Delta RBD-LNP in half dose. Interestingly, both Omicron-specific and Hybrid RBD-LNP elicited extremely high titer of neutralizing antibody against Omicron itself, but few to none neutralizing antibody against other SARS-CoV-2 variants. The bivalent RBD-LNP, on the other hand, generated antibody with broadly neutralizing activity against the wild-type virus and all variants. Surprisingly, similar cross-protection was also shown by the Delta-specific RBD-LNP. Taken together, our data demonstrated that Omicron-specific mRNA vaccine can induce potent neutralizing antibody response against Omicron, but the inclusion of epitopes from other variants may be required for eliciting cross-protection. This study would lay a foundation for rational development of the next generation vaccines against SARS-CoV-2 VOCs.

## Introduction

Since the COVID-19 pandemic occurred in late 2019, vaccine has been regarded as a major approach to combat the disease. Currently, global research and clinical efforts have pushed several United States Food and Drug Administration (U.S. FDA)-approved COVID-19 vaccines for clinical use ^1^. However, the pandemic is still far from over due to the constant emergence of new SARS-CoV-2 variants of concern (VOC) ^2^. Among the earlier identified VOCs, B.1.351 (Beta) exhibited the greatest immune escape against convalescent sera obtained from COVID-19 patients or vaccinated individuals ^3,4^. The B. 1.617.2 (Delta) variant that emerged in early December, 2020, quickly outpaced all other circulating isolates and showed a significant reduction in vaccine effectiveness. Importantly, acquisition of favorable mutations in Delta strain enhances transmissibility among individuals and leads to more severe outcomes ^5,6^. In late November 2021, the B.1.1.529 (Omicron) variant was first discovered and rapidly spread globally. This variant contains novel genomic sequence changes different from any of the previously defined ancestral or VOC isolates of SARS-CoV-2, including 37 mutations in the spike protein, 15 of which are located in the RBD ^7^. Recent studies have shown that an increase in the number and complexity of spike mutations leads to inability of therapeutic monoclonal antibodies against Omicron strain ^8^. Furthermore, constellation mutations render Omicron more antigenically distant from ancestral viruses or other VOCs, leading to reduced antibody neutralizing activity from vaccination or natural infection ^7,9^. Although the disease symptoms induced by Omicron variant are milder than that of Delta ^10^, higher transmission rates may inevitably lead to an increase in case numbers and pose a threat on the public health and economics of society. Therefore, there is an urgent need to develop a new generation of vaccines to prevent from VOCs pandemic.

In this study, we developed monovalent receptor-binding domain (RBD)-based mRNA vaccines targeting on two currently major predominant VOCs, Omicron and Delta. We also tested the concept of bivalent vaccines containing both Delta and Omicron RBD, and a Hybrid vaccine, which combined the mutation sites of Delta and Omicron in single RBD construct since multivalent vaccines containing various SARS-CoV-2 VOC antigens are recommended to effectively control the spread of SARS-CoV-2 variants according to the recommendations of the WHO Technical Advisory Group on COVID-19 Vaccine Components (TAG-CO-VAC). Using pseudovirus neutralization assays, we found that serum samples from the Omicron vaccinated mice can effectively neutralize Omicron, but not the wild-type (D614G), or other VOCs (Beta and Delta) of SARS-CoV-2. In contrast, the Omicron/Delta bivalent mRNA vaccine elicited broadly cross-reactive neutralizing antibodies, effectively neutralizing Omicron and other VOCs. Taken together, our data demonstrate that a new generation of multivalent COVID-19 mRNA vaccine is a viable approach to prevent infection from ancestral or VOCs of SARS-CoV-2.

## Results

### Design and encapsulation of mRNA encoding variant RBD

To cope with the emergence of new variants, 4 different mRNA vaccines were designed to encode the SARS-CoV-2 spike receptor-binding domain (RBD) region of wild-type (WT, Wuhan strain), Delta, Omicron, and Omicron with additional L452R mutation (named Hybrid), respectively. The RNA constructs with mutation sites were summarized in figure 1A. The *in vitro* transcription reaction was used to synthesize mRNA and the fragment analysis was conducted to analyze RNA integrity. Four synthesized RNA had expected length (around 1000nt) and showed great integrity with 93% or 94% of intact RNA and only limited amounts of degraded transcripts (Fig 1B and Sup 1). The mRNA was then transfected to 293T cells for RBD expression examination. Two days post transfection, supernatants were collected and cocultured with 293T cells that stably expressed human angiotensin-converting enzyme 2 (293T-hACE2) to conduct binding assay. The bound RBD was then detected by polyclonal anti-RBD antibodies. All WT, Delta, Omicron, and Hybrid RBD mRNA efficiently expressed RBD with around 99% of cells stained positive in each construct. Also, we assessed the ability of expressed RBD to bind mouse ACE2 as previous study had shown that RBD of Omicron variant gained the ability to bind mouse ACE2 ^7^. In this case, 3T3 cells that stably expressed mouse ACE2 were used. In contrast to WT and Delta RBD, which showed no binding capacity against mouse ACE2, RBD from Omicron and Hybrid mRNA transfected supernatants efficiently bound to mouse ACE2 (Fig 1C). The synthesized mRNAs were then packaged into lipid nanoparticle (LNP) to get WT, Delta, Omicron, and Hybrid RBD-LNP vaccines. In addition to these 4 constructs, we also formulated half dose of both Delta and Omicron mRNAs into the same LNP at 1:1 ratio to get bivalent RBD-LNP vaccine (Fig 1D). Dynamic light scattering (DLS) measurement showed that the average size of these LNP ranged between 86 nm and 99 nm with a narrow distribution (pdI around 0.121 to 0.147). The zeta potential was about 6 - 9 mV (Sup 2). We also examined the RBD expression capacity of these RBD-LNP by transfection into 293T cells. Two days post transfection, supernatants were collected and cocultured with 293T-hACE2 cells to conduct binding assay. All 5 RBD-LNP vaccine efficiently expressed RBD with around 99% of cells stained positive in each group (Fig 1D).

**Fig. 1.**
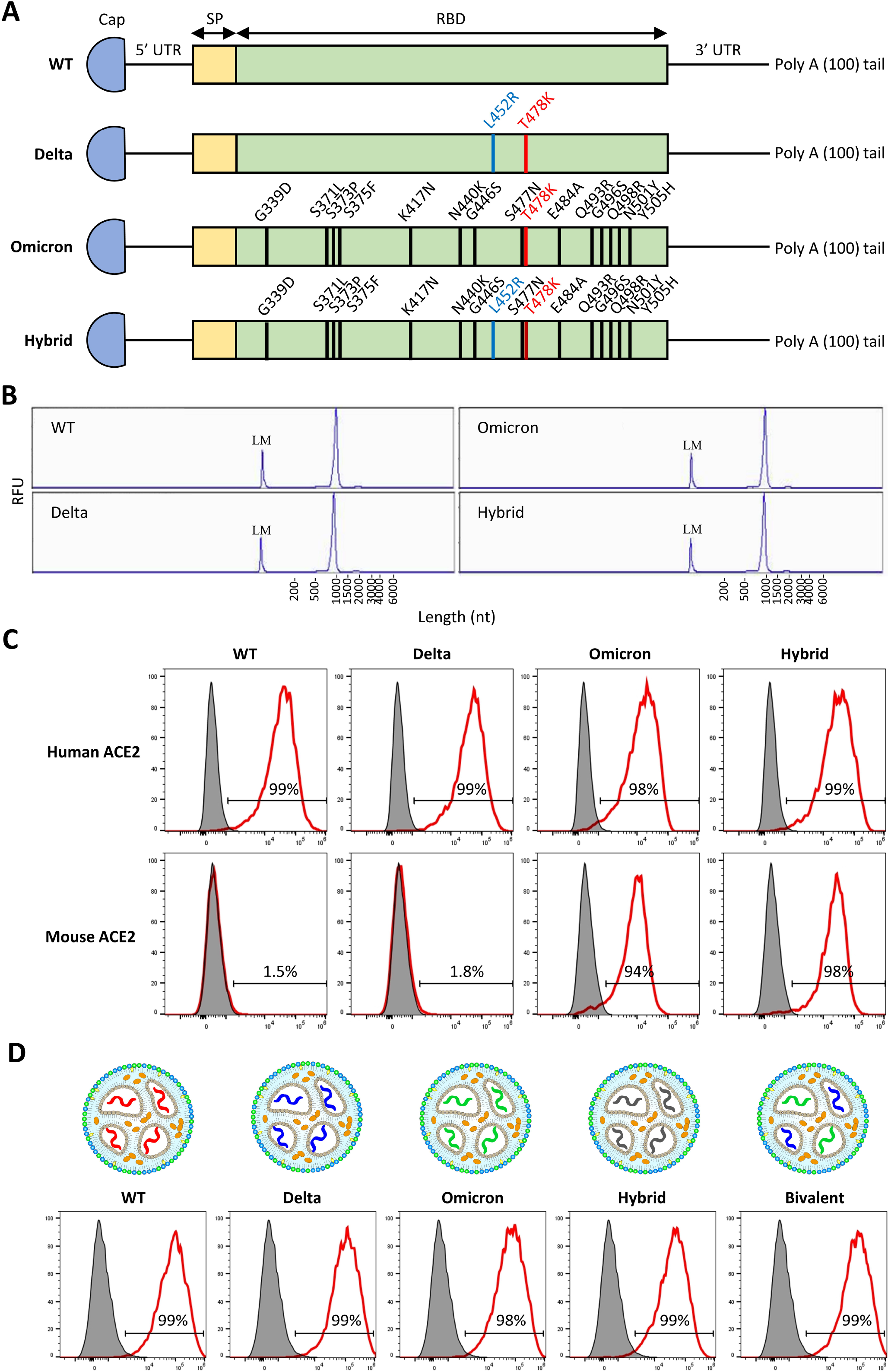
RBD mRNA constructs and RBD-LNP vaccines. (A) Mutation sites of wildtype (WT), Delta, Omicron, and Hybrid RBD mRNA constructs. UTR, untranslated region. SP, signal peptide. (B) Fragment analysis of RNA identity and integrity of in vitro transcribed WT, Delta, Omicron, and Hybrid RBD mRNA. LM, lower marker. (C) FACS analysis of RBD binding against human ACE2 or mouse ACE2 of indicated RBD mRNA in transfected cell supernatants. (D) Schematic illustration of WT, Delta, Omicron, Hybrid, and Delta/Omicron bivalent RBD-LNP and FACS analysis of RBD expression of indicated RBD-LNP in transfected cell supernatants.

### Immunogenicity of various RBD-LNP vaccine

Next, we assessed the ability of various RBD-LNP vaccine to generate neutralizing antibody responses. Group of BALB/c mice were immunized intramuscularly with WT, Delta, Omicron, Hybrid, and bivalent RBD-LNP for 2 times at 2-week interval. Serum samples were collected 1 week post second immunization and subjected to SARS-CoV-2 pseudovirus neutralization assay. The neutralization curves were shown at the left panel and the 50% neutralizing titer (NT50) values were summarized at right (Fig 2). Sera that collected from mice immunized with WT vaccine showed great neutralizing capacity against D614G, Beta, and Delta variants of SARS-CoV-2 pseudovirus with mean NT50 about 6,400, 3,000, and 4,000, respectively. However, the neutralization capacity was significant lower against Omicron variant with about 8-fold decline (mean NT50 about 503). On the contrary, the Omicron RBD-LNP vaccine showed extremely high neutralizing antibodies against Omicron variant with mean NT50 about 18,600 but failed to neutralize other tested SARS-CoV-2 variants. The bivalent RBD-LNP, which contained half dose of both Delta RBD mRNA and Omicron RBD mRNA, generated high titer of neutralizing antibodies against all D614G, Beta, Delta, and Omicron variants with mean NT50 about 8,600, 1,700, 10,500, and 4,000, respectively. Surprisingly, Delta RBD-LNP also showed high titer of neutralizing antibodies against all D614G, Beta, Delta, and Omicron variants with mean NT50 about 11,700, 2,200, 17,800, and 4,000, respectively. The Hybrid RBD-LNP showed extremely high titer of neutralizing antibodies against Omicron variant with mean NT50 about 21,100 but low neutralizing antibody titer against other tested variants (mean NT50 about 300 against D614G, 500 against Beta, and 400 against Delta variants). Taken together, we found that Omicron RBD-LNP vaccine could generate potent neutralizing antibody responses against Omicron variant but had no neutralizing capacity against other variants. By contrast, the bivalent RBD-LNP vaccine or the Delta RBD-LNP vaccine could generate broad neutralizing antibody responses against ancestral SARS-CoV-2 virus and variants.

**Fig. 2.**
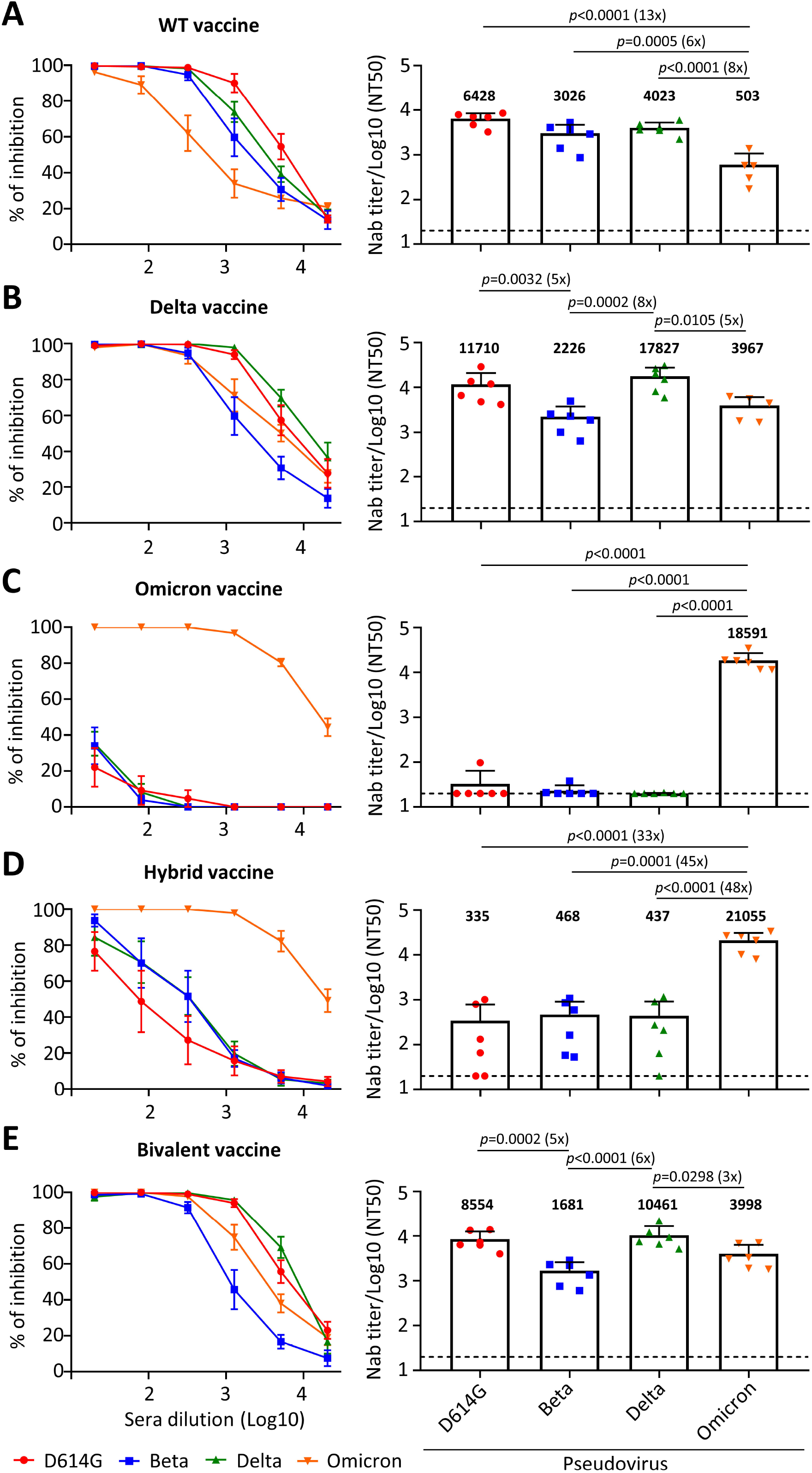
Neutralization capacity of various RBD-LNP vaccine immunized mouse sera against D614G, Beta, Delta, and Omicron SARS-CoV-2 pseudovirus variants. (A-E) Neutralization curves (left panel) and summarized NT50 values (right panel) of vaccinated mouse sera against pseudotyped SARS-CoV-2 and the variants. Mean NT50 titers are shown above each column. Dashed lines indicate the limit of detection. NT50, 50% neutralization titer. Data are presented as mean ± SD. *p* value were calculated by one-way ANOVA.

## Discussion

Recently, several studies analyzing the sera from vaccinated or convalescent subjects revealed that the major antigenic shift of Omicron variant lead to immune evasion ^7,11–13^. By using vaccinated mouse model which provided an identical genetic background and immune profile, we fairly assessed the neutralizing antibody response induced by various RBD mRNA-LNP vaccines. Our data showed that WT vaccine can induce high neutralization titers against D614G, Beta, and Delta variants (Fig 2A), but only caused a marginal effect (7.8% of D614G) to Omicron variant (Fig 2A). The loss of WT vaccine-mediated immunity against Omicron may be due to the loss of epitopes critical for neutralizing antibody recognition since it has been reported that mutations on Omicron variant such as K417N, G446S, E484A, and Q493R impaired a large panel of monoclonal antibodies under commercial development ^7,8,14,15^. In addition, a cross-variant protection of WT vaccine against Omicron still existed (Fig 2A), indicating that certain conserved epitopes shared by WT and Omicron may confer neutralization effect, despite low immunogenic and limited protection to Omicron. However, we do not know whether Omicron-specific vaccine can induce such an immune response in people who have been immunized with vaccines based on ancestral SARS-CoV-2 strain. To answer this question, immunoanalysis by using mice received heterologous WT/Omicron prime-boost vaccination is needed.

To efficiently prevent Omicron pandemic, we generated an Omicron-specific vaccine which can elicit extremely high neutralizing antibody titers against Omicron itself but failed to neutralize other SARS-CoV-2 variants (Fig 2C). However, at present, Delta is still another dominant variant associated with more severe illness, making up 28% of all cases as of 22nd Jan (COVID-19 Weekly Epidemiological Update, 75th edition). To simultaneously prevent spread of Delta and Omicron, we designed a bivalent vaccine which contained both RBD-LNPs in half dose. Our data showed that combinatorial vaccination generated broadly neutralizing activity (Fig 2E). Although the neutralizing activity elicited by bivalent vaccine was lower than that generated by the full dose Omicron vaccine (Fig 2C) or Hybrid vaccine (Fig 2D), perhaps because of Omicron RBD dose halved, bivalent vaccine is still a potent strategy to increase the breadth and potency of vaccine. In the future, different Delta/Omicron RBD-LNP ratios can be tested for improvement of vaccine effectiveness. To our surprise, monovalent Delta RBD-LNP also showed cross-strains immunity against Omicron (Fig 2B). Since a recent study reported that Delta virus infection also induced a cross-variant neutralization of Omicron ^16^, it is reasonable to design next generation vaccines based on the Delta RBD sequence. Taken together, our data demonstrated that Omicron-specific mRNA vaccine induced potent neutralizing antibody response against Omicron but not other SARS-CoV-2 variants and lay the foundation for rational development of next generation vaccines against SARS-CoV-2 VOCs.

## Materials and Methods

### Ethics statement

All mouse works were conducted in accordance with the “Guideline for the Care and Use of Laboratory Animals” as defined by the Council of Agriculture, Taiwan and was approved by the Institutional Animal Care and Use Committee of Academia Sinica (protocol ID: 20-05-1471).

### Animals

BALB/c mice were purchased from the National Laboratory Animal Center (Taipei, Taiwan) and maintained in a specific pathogen-free environment in the animal facilities of the Institute of Biomedical Sciences, Academia Sinica. All experimental procedures were reviewed and approved by the Animal Care and Use Committee of Academia Sinica.

### Generation of modified mRNA

DNA templates, which incorporated 5’ untranslated regions (UTR) (GGGAAAUAAGAGAGAAAAGAAGAGUAAGAAGAAAUAUAAGAGCCACC), signal peptide sequences from Igκ (ATGGAGACAGACACACTCCTGCTATGGGTACTGCTGCTCTGGGTTCCAGGTTCCACCGGTGAC), codon optimized wildtype (Wuhan-Hu-1, GenBank YP_009724390.1), Delta, Omicron, and Omicron with additional L452R (Hybrid) RBD sequence, 3’ UTR (UGAUAAUAGGCUGGAGCCUCGGUGGCCAUGCUUCUUGCCCCUUGGGCCUCCCCCCAGCCCCUCCUCCCCUUCCUGCACCCGUACCCCCGUGGUCUUUGAAUAAAGUCUGA), and a poly-A tail were constructed. Before subjected to the *in vitro* transcription reaction to synthesize mRNA with T7 RNA polymerase (NEB, MA,USA), the DNA template was linearized with EcoRV (NEB, MA,USA). The *in vitro* transcription reaction included CleanCap^®^Reagent AG (3’ OMe) (Trilink, CA, USA) for co-transcriptional capping of mRNA and complete replacement of uridine by N1-methyl-pseudouridine (Trilink, CA, USA). The mRNA was purified by LiCl (Invitrogen, MA, USA) precipitation and dsRNA was depleted by cellulose (Sigma-Aldrich, MA, USA). Purified RNA was kept frozen at −80 °C until further use.

### Fragment analysis

RNA integrity was analyzed by fragment analysis following manufactural protocol (Agilent, CA, USA). Briefly, mRNA was diluted to 2 ng/μl and mixed with diluent marker. RNA samples and ladder were denatured at 70°C for 2 minutes and kept on ice before use. The percentage of RNA integrity was quantified by smear analysis of ProSize Data Analysis Software (Agilent, CA, USA).

### Preparation of RBD-LNP

The RBD mRNAs were added to an ethanol solution containing a lipid mixture of cationic lipid, DMG-PEG2000 (MedChemExpress, NJ, USA), 1,2-distearoyl-sn-glycero-3-phosphocholine (DSPC) (Avanti, NY, USA), and cholesterol (Sigma, MA, USA). The weight ratio of the mRNA and the lipid in the ethanol solution was 3: 1. The mixtures were subjected to the NanoAssemblr IGNITETM NxGen Cartridges (Precision NanoSystems, BC, Canada) to produce mRNA-LNP composition, followed by treatments of dialysis against Dulbecco’s phosphate buffered saline (DPBS) (Gibco, MA, USA). The size and zeta potential of the RBD-LNP were measured by Zetasizer Nano ZS (Malvern Panalytical Ltd., Malvern, WR, U.K.).

### RBD expression and binding assay

The variant-specific RBD mRNA was transfected into 293T cells via lipofectamine (Invitrogen, MA, USA) and the variant-specific RBD-LNP was transfected by directly added. Cell supernatants were collected 2 days post transfection. To test the ability of RBD binding to human ACE2 or mouse ACE2, 293T-hACE2 or 3T3-mACE2 cells were harvested and aliquoted into FACS tubes at 5×10^5^ cells/tube. The cells were washed with staining buffer (DPBS+1% BCS) and then incubated in 100μl of transfected cell supernatant at 4°C for 1 hour. After washing, the cells were incubated with anti-RBD polyclonal antibody (1μg/tube) at 4°C for 30 minutes. The cells were then washed two times, followed by 30-minute incubation with PE-goat-anti-mouse IgG (H+L) antibody (Jackson ImmunoResearch, PA, USA) at 4°C. The cells were washed twice and resuspended in 300 μl of staining buffer containing 7-AAD (Biolegend, CA, USA) for flow cytometry analysis (Thermo Fisher Attune NxT - 14 color analyzer, Thermo Fisher Attune NxT software v2.2, FlowJo 10.6.1).

### Immunization

Group of BALB/c mice were respectively immunized intramuscularly with two doses of WT (10 μg per dose), Delta (10 μg per dose), Omicron (10 μg per dose), Hybrid (10 μg per dose), and bivalent (5 μg of both Delta and Omicron RBD mRNA per dose) with an interval of 2 weeks. The serum samples were collected from the mice 1 week post last immunization.

### SARS-CoV-2 pseudovirus neutralization assay

293T cells that stably expressed human ACE2 (293T-hACE2) and lentiviral-based pseudotyped SARS-CoV-2 viruses were provided by National RNAi Core Facility (Academia Sinica, Taiwan). One day before neutralization assay, 293T-hACE2 cells were seeded into 96-well black plate (Perkin Elmer, MA, USA) at a density of 1 x 10^4^ cells per well at 37°C. Mouse sera were inactivated at 56°C for 30 minutes and performed four-fold serial dilutions with culture medium before incubation with indicated SARS-CoV-2 pseudovirus for an hour. The mixtures were then added to pre-seeded 293T-hACE2 cells and incubated for 3 days. Luciferase activity was measured by Luciferase Assay kit (Promega, WI, USA). The 50% neutralization titer (NT50) was calculated by nonlinear regression using Prism software version 8.1.0 (GraphPad Software Inc.).

### Statistical analysis

Results are presented as the mean ± standard deviation (SD). Differences between experimental groups of animals were analyzed by one-way ANOVA with Tukey’s comparison. *p* < 0.05 was considered as statistically significant.

## Acknowledgments

We thank all study participants who devoted time to our research and Dr. James C. Liao for helpful discussions and advice. We thank the Academia Sinica SPF Animal Facility (AS-CFII-111-204) for providing animal support, Flow Cytometry Core Facility of Institute of Biomedical Sciences, Academia Sinica (AS-CFII108-113) for supplying flow cytometry instrumentation, DNA Sequencing Core Facility of Institute of Biomedical Sciences, Academia Sinica (AS-CFII-111-211) for sequencing support, National RNAi Core Facility at Academia Sinica for providing 293T-hACE2 cell line and SARS-CoV-2 pseudovirus. This work was supported by Academia Sinica Biomedical Translation Research Center grant AS-KPQ-110-EIMD and AS-KPQ-111-KNT.

## Author contributions

M.H.T., I.J.L., P.Y.W. conceived and designed the project. M.H.T. provided overall instruction for the study and supervised the project. I.J.L coordinated the experiments. I.J.L., Y.H.L., Y.S.L., H.F.C., and T.Y.C. performed animal experiments. S.C.T. was responsible for construction of mRNA expression vectors. Y.W.C. and C.C.L. performed RNA in vitro transcription. C.M.C. and M.K. packaged mRNA-LNPs. Y.C.C provided VSV-based pseudotyped SARS-CoV-2. H.T.L. and W.Y.C. produced monoclonal antibodies for ELISA. I.J.L., Y.H.L., P.Y.W., S.I.T., C.W.C., C.H.H., and C.Y.C. conducted cell-based binding assay and pseudovirus neutralization assay. I.J.L., M.H.T., and P.Y.W. analyzed the data. H.C.W., C.P.C., and C.C.L. provided consultations. C.P.C. and Y.J.L. wrote the original draft with input from the team. I.H.W., W.C.L, and M.H.T. reviewed and edited the manuscript.

*Sup. 1 Summary bar plot of RNA integrity*

(A) Fragment analysis of RNA integrity of in vitro transcribed WT, Delta, Omicron, and Hybrid RBD mRNA. H.M.W, high molecular weight. The percentages are shown above each column.

*Sup. 2 Basic characteristics of various RBD-LNP*

(A) The summary table of size, polydispersity index (pdI), and zeta potential of indicated RBD-LNP. Data are presented as mean ± SD.

## References

1 Garcia-Montero, C. et al. An Updated Review of SARS-CoV-2 Vaccines and the Importance of Effective Vaccination Programs in Pandemic Times. Vaccines 9, doi:10.3390/vaccines9050433 (2021).

2 Freer, G., Lai, M., Quaranta, P., Spezia, P. G. & Pistello, M. Evolution of viruses and the emergence of SARS-CoV-2 variants. New Microbiol 44 (2021).

3 Cele, S. et al. Escape of SARS-CoV-2 501Y.V2 from neutralization by convalescent plasma. Nature 593, 142–146, doi:10.1038/s41586-021-03471-w (2021).

4 Collier, D. A. et al. Sensitivity of SARS-CoV-2 B.1.1.7 to mRNA vaccine-elicited antibodies. Nature 593, 136–141, doi:10.1038/s41586-021-03412-7 (2021).

5 Lopez Bernal, J. et al. Effectiveness of Covid-19 Vaccines against the B.1.617.2 (Delta) Variant. N Engl J Med 385, 585–594, doi:10.1056/NEJMoa2108891 (2021).

6 Sheikh, A. et al. SARS-CoV-2 Delta VOC in Scotland: demographics, risk of hospital admission, and vaccine effectiveness. Lancet 397, 2461–2462, doi:10.1016/S0140-6736(21)01358-1 (2021).

7 Cameroni, E. et al. Broadly neutralizing antibodies overcome SARS-CoV-2 Omicron antigenic shift. Nature, doi:10.1038/s41586-021-04386-2 (2021).

8 VanBlargan, L. A. et al. An infectious SARS-CoV-2 B.1.1.529 Omicron virus escapes neutralization by therapeutic monoclonal antibodies. Nat Med, doi:10.1038/s41591-021-01678-y (2022).

9 Cele, S. et al. SARS-CoV-2 Omicron has extensive but incomplete escape of Pfizer BNT162b2 elicited neutralization and requires ACE2 for infection. medRxiv, doi:10.1101/2021.12.08.21267417 (2021).

10 Dejnirattisai, W. et al. Reduced neutralisation of SARS-CoV-2 omicron B. 1.1.529 variant by post-immunisation serum. Lancet 399, 234–236, doi:10.1016/S0140-6736(21)02844-0 (2022).

11 Carreno, J. M. et al. Activity of convalescent and vaccine serum against SARS-CoV-2 Omicron. Nature, doi:10.1038/s41586-022-04399-5 (2021).

12 Sievers, B. L. et al. Antibodies elicited by SARS-CoV-2 infection or mRNA vaccines have reduced neutralizing activity against Beta and Omicron pseudoviruses. Sci Transl Med, eabn7842, doi:10.1126/scitranslmed.abn7842 (2022).

13 Cele, S. et al. Omicron extensively but incompletely escapes Pfizer BNT162b2 neutralization. Nature, doi:10.1038/s41586-021-04387-1 (2021).

14 Dejnirattisai, W. et al. SARS-CoV-2 Omicron-B.1.1.529 leads to widespread escape from neutralizing antibody responses. Cell, doi:10.1016/j.cell.2021.12.046 (2022).

15 Cao, Y. et al. Omicron escapes the majority of existing SARS-CoV-2 neutralizing antibodies. Nature, doi:10.1038/s41586-021-04385-3 (2021).

16 Suryawanshi, R. K. et al. Limited cross-variant immunity after infection with the SARS-CoV-2 Omicron variant without vaccination. medRxiv, doi:10.1101/2022.01.13.22269243 (2022).

