## Supplementary figures and images for "Omicron-specific mRNA vaccine induced potent neutralizing antibody against Omicron but not other SARS-CoV-2 variants"

### Supplementary Figure 1

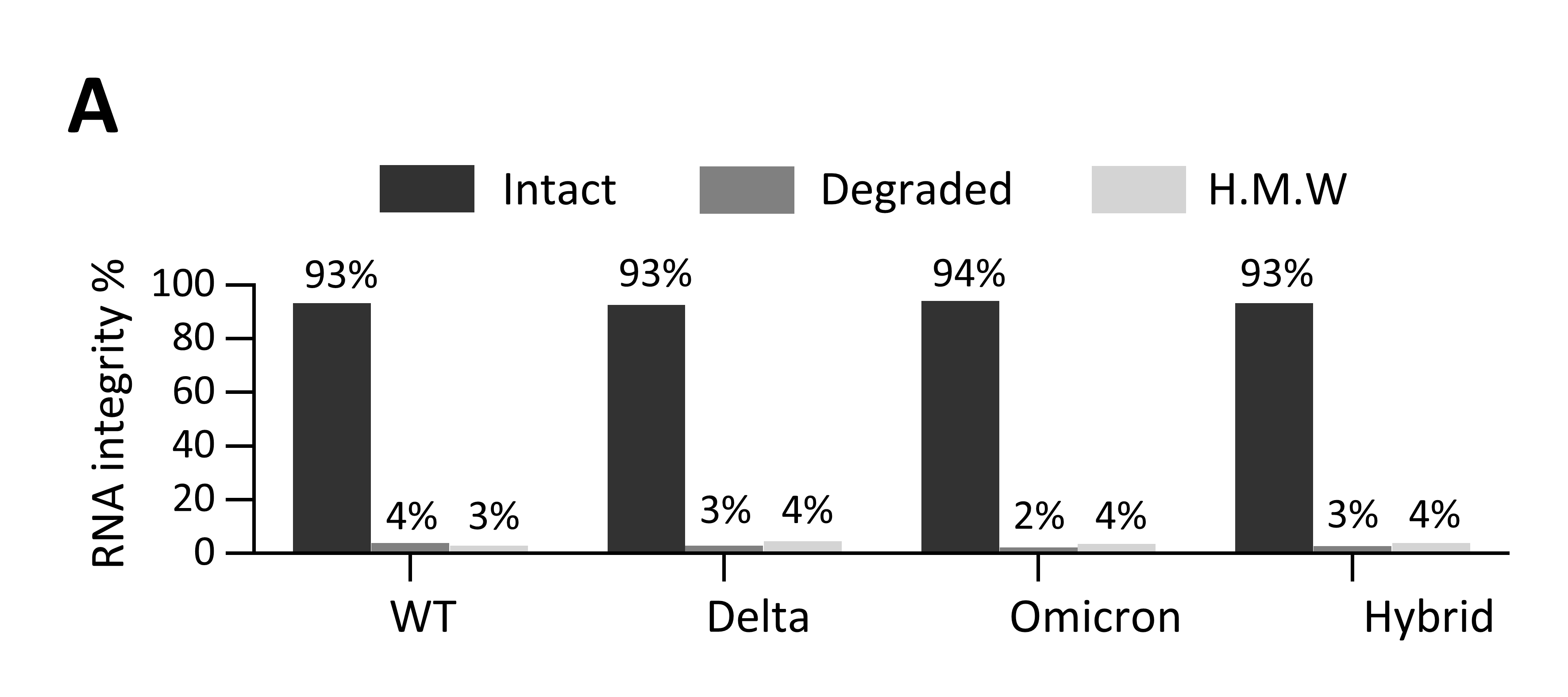

### Supplementary Figure 2

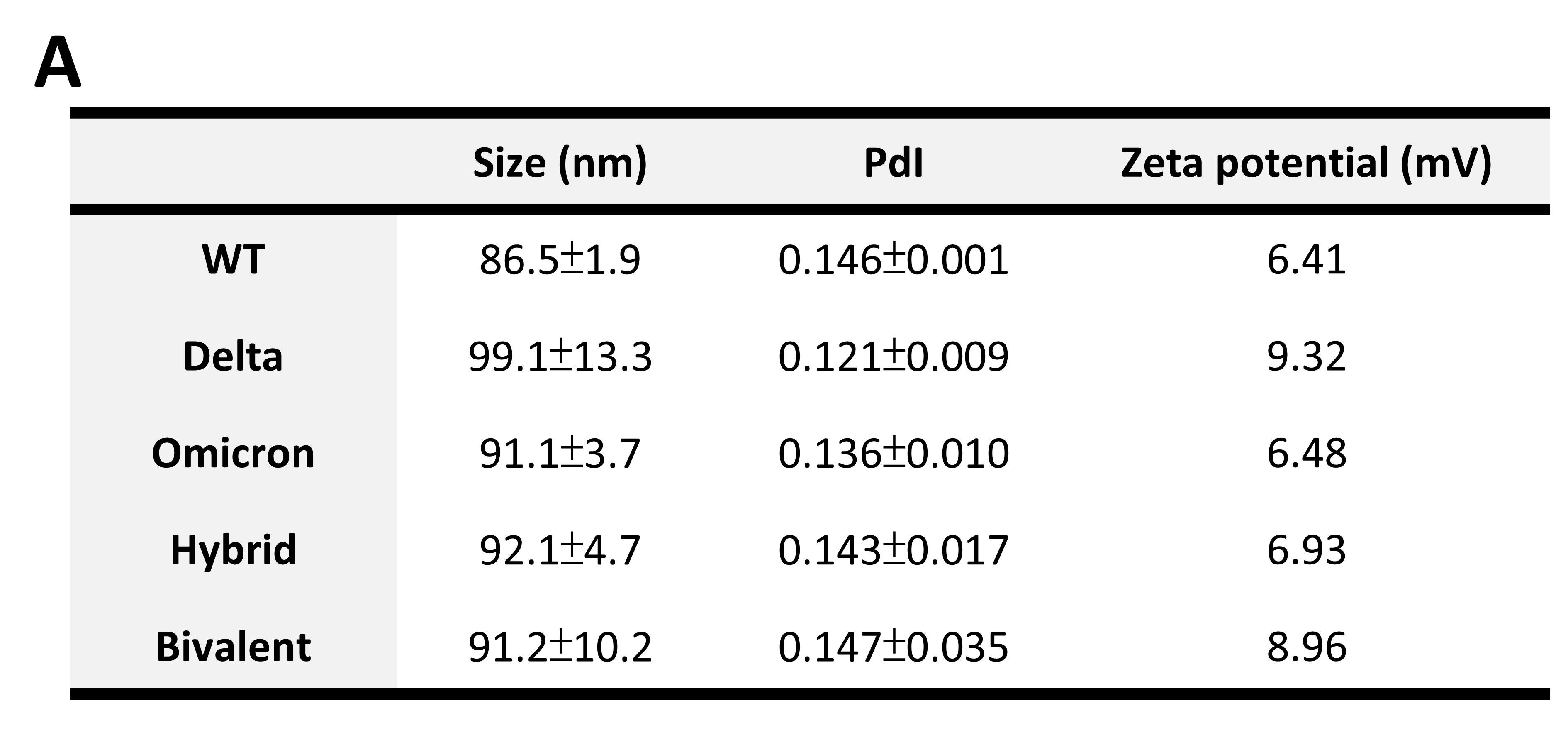
